# From Metabolomics to Function: Ranking Plant Stem Cell Metabolomes for Use in Health and Cosmetics

**DOI:** 10.64898/2026.03.17.711179

**Authors:** Assaf Zemach, Mikhail R. Plaza, Bong Seop Lee, Leo Little Dog, Efrain Santiago-Rodriguez, Dior Simmons, Melissa Palomares, Dodanim Talavera-Adame, Nathan Newman

## Abstract

**Background:** Plants produce diverse metabolites with potential benefits for human health. However, the metabolomes of plant callus cultures—cell cultures analogous to stem cells—remain poorly characterized in terms of their functional relevance.

**Methods:** We profiled the metabolomes of six plant calli: *Acacia concinna* (Shikakai), *Daucus carota* (carrot), *Hibiscus sabdariffa* (hibiscus), *Linum usitatissimum* (flax), O*cimum sanctum* (tulsi), and the *Nicotiana tabacum* Bright-Yellow 2 (BY-2) cell line. To facilitate functional interpretation, we developed Metabolite2Function (M2F), a pipeline that annotates metabolites with biological functions using scientific literature and large language modeling.

**Results:** Untargeted metabolomics identified 177 metabolites, revealing clustering patterns independent of genetic relationships, culture age, or growth rate. Tulsi and carrot calli exhibited enrichment in metabolites relative to the tobacco reference line, whereas flax and hibiscus were comparatively depleted. Most metabolites varied across at least four calli, and 10% were unique to a single species. Using M2F, we annotated 87 metabolites with beneficial activities, including antioxidant, anti-glycation, anti-inflammatory, and anti-senescence functions, as well as skin-related effects such as collagen production and brightening. Notably, antioxidant and anti-senescence metabolite levels correlated with corresponding biological activities in human cells.

**Conclusions:** Plant callus cultures generate distinct and functionally diverse bioactive metabolomes. M2F provides a scalable framework for systematic functional annotation relevant to human health and cosmetic applications.

## 1. Introduction

Plants are known to contain numerous bioactive compounds that are beneficial to human health and have been used for medicinal and cosmetic purposes [1,2]. Beyond serving as a source of energy, plant compounds, known as phytochemicals, play important roles in human development, health, and personal care [3]. In some cases, phytochemicals are essential for normal growth and development, contribute to the prevention and treatment of various diseases and conditions, and enhance overall quality of life [4,5].

Plants produce diverse sets of compounds within their various organs and tissues, making it essential to select the appropriate plant organ or tissue when extracting phytochemicals [6]. Flowers and fruits are particularly rich in phytochemicals, as these tissues evolved to attract and reward animals [7]. For instance, resveratrol is concentrated in grape skins, while cannabinoids are abundant in cannabis flowers [8,9]. In contrast, green tissues such as leaves, which are photosynthetically active and metabolically dynamic, are enriched in antioxidants and bioactive compounds, including epigallocatechin-3-gallate from the tea plant (*Camellia sinensis*) [10]. Other organs also provide valuable compounds, such as curcumin and β-carotene found in the roots of turmeric (*Curcuma longa*) and carrot (*Daucus carota*), respectively [11,12].

Since the domestication of plants approximately 10,000 years ago, humans have sought to industrialize their cultivation for food, health, and personal care applications. In the past century, and especially in recent decades, plant tissue culture techniques have been developed to optimize the production of bioactive compounds [13,14]. By strategically applying plant growth regulators, cells can be induced to transdifferentiate into tissue cultures with advantageous industrial properties, including stability, consistency, and rapid growth [15]. Tissue cultures can be established either on semi-solid media (e.g., agar), where they typically form callus tissue, or in liquid suspension.

Callus, which naturally arises in plants in response to stimuli such as injury or infection, possesses the capacity for self-renewal and retains the potential to differentiate into various cell types, paralleling stem cell–like properties [16–19]. Importantly, the process of trans-differentiation alters the developmental stage of the cells, resulting in chemical compositions distinct from those of the original tissues. Although the phytochemical profiles of intact plant organs and extracts are well characterized, plant tissue cultures remain comparatively underexplored, despite emerging evidence supporting the potential of plant calli as sources of beneficial natural products [20,21]. This study highlights the need for further exploration into the phytochemical profiles of plant tissue cultures, particularly in terms of their bioactive properties. Such investigations could unlock novel applications in pharmaceuticals and nutraceuticals, providing consistent and sustainable sources of bioactive compounds, and improving their therapeutic efficacy.

Within the past few years, the field of Natural Language Processing has grown substantially with the advent of large language models (LLMs). One of these LLM models, OpenAI’s GPT, has grown in popularity for its extensive language comprehension and generative abilities, and its ability to automate tasks. Text annotation is traditionally a time-intensive process, and researchers have used GPT to automate large sets of text data [22]. GPT has been reported to be more accurate than crowd-sourced and individually trained annotators, resulting in time and cost-savings. Recently, GPT was demonstrated to have outperformed physicians [23] and other AI tools [24] in literature searches for systematic reviews.

Despite the growing interest in plant-derived compounds [25], the absence of comprehensive and accessible metabolome ontology databases is a significant bottleneck in our ability to systematically characterize the functional roles of these metabolites. While major databases have been created for humans, including the Human Metabolome Database (HMDB, https://www.hmdb.ca/), their coverage on metabolite function and biological role is limited. Such databases often lack detailed annotations, linking metabolites to biological processes, mechanisms of action, or potential health benefits. Additionally, there are very few resources describing plant metabolites and their specific functions and bioactivities. This gap presents a significant barrier to integrating metabolomic data with function, especially with metabolites from understudied plant species. To address this limitation, we developed an automated metabolomics ontology pipeline that is based on text annotation of scientific publications by the GPT model.

Metabolomics has emerged as a powerful tool to characterize the metabolite composition of plant calli. Profiling these metabolites is critical to identifying compounds with medicinal and cosmetic potential. This study focuses on understanding the metabolites that are found within six different plant calli and their prospective potential benefit in humans using an untargeted metabolomics approach and testing their effects on human cells culture. Our findings may contribute to the development of innovative, plant-based products that harness the unique metabolite signatures of plant calli.

## 2. Materials and Methods

### 2.1. Cell Cultures

Shikakai, carrot, hibiscus, flax, and tulsi plant cell cultures (calli) were established from *Acacia concinna, Daucus carota, Hibiscus sabdariffa, Linum usitatissimum*, and *Ocimum sanctum* seedlings using plant growth hormones, respectively. Tobacco Bright Yellow-2 (BY-2) callus culture was obtained from Leibniz Institute DSMZ. Plant calli were cultivated on 0.8% agar Murashige and Skoog (MS) or Gamborg B5 media, supplemented with 3% sucrose and plant growth hormones. Cultures were kept at room temperature 20°C to 25°C either in the dark or under 16:8 light:dark cycle and were sub-cultured every three to four weeks.

### 2.2. UV absorbance

Plant calli were extracted in 1:1 methanol:water solution, using a homogenizer, and the resuspended-filtered portion was subjected to UV absorbance in 10 mm cuvettes and the DeNovix DS11 FX+.

### 2.3. Dry matter content

Dry matter content was determined by gravimetric moisture analysis. Briefly, 1 g of fresh callus tissue from each plant callus line was weighed and placed onto an aluminum sample pan, with three biological replicates analyzed per plant. Samples were gently tapped to flatten the callus and ensure uniform heating. Moisture content was measured using a Shimadzu Unibloc Moisture Analyzer (MOC63u), and dry matter content was calculated from the weight loss during drying.

### 2.4. Metabolomic analysis

Media derived from the plant cell cultures were sent to the Carver Metabolomics Core, University of Illinois Urbana-Champaign Roy. J. Carver Biotechnology Center (Urbana, IL) for untargeted metabolomics to profile compounds found in each sample. Processed samples were dried, reconstituted in 100 µL methanol:water, and had 5 µL injected onto the instrument for analysis. Samples were analyzed using a Dionex Ultimate 3000 series UHPLC system (Thermo Scientific) with a Q-Exactive MS system (Thermo Scientific), as described previously [26]. Samples were analyzed with a reversed phase liquid chromatography (RPLC) performed using a Waters Acquity ethylene-bridged hybrid (BEH) C18 column (100 mm × 2.1 mm; 1.7 μm) column maintained at 25°C with a flow rate of 0.3 - 0.4 mL/min (Waters Corp). The mobile phases consisted of (A) water including 0.1% formic acid and solvent (B) acetonitrile including 0.1% formic acid. Spectra were acquired in positive and negative ionization mode.

All the LC-MS raw data files were performed using MS-DIAL ver.4.90 software for data collection, peak detection, alignment, adduct, and identification [27]. The detailed parameter setting was as follows: MS1 tolerance, 0.005 Da; MS2 tolerance, 0.01 Da; minimum peak height, 10,000 amplitude; mass slice width, 0.05 Da; smoothing method, linear weighted moving average; smoothing level, 3 scans; minimum peak width, 5 scans. [M-H]-, [2M-H]- and [M+H]+, [2M+H]+, [M+NH4]+, [M+Na]+, [M+2H]2+ were included in adduct ion setting for negative and positive mode, respectively. Compounds were annotated by m/z, MS/MS spectra, and retention time against an in-house library produced using chemical standards. In addition, they were annotated by m/z and MS/MS spectra using the public libraries’ MassBank of North America (MoNA), MassBank Europe (MassBank EU), and Global Natural Products Social Molecular Networking (GNPS), as well as the commercial library National Institute of Technology 20 (NIST20). Internal standards were monitored for retention time and intensity, and principal component analysis (PCA) was used for multivariate statistics and visualization, specifically for outlier detection.

From the MS-DIAL results file, all detected features/metabolites were removed if (sample max)/(blank average) <10. Known (positively identified/annotated) features/metabolites were evaluated for MSI level 1 matches [28]. Next, positive and negative mode data were combined and replicated identifications removed by initially retaining features with MSI level 1 match. Due to the MS-DIAL software not evaluating MS/MS for metabolites with both m/z and retention time matches, manual evaluation of spectra was performed to confirm level 1 identifications. Manual inspections of spectra MSI level 2 were conducted in certain circumstances flagged by specific parameters that required further investigation of identifications. Remaining replicate features without MSI level 1 match were filtered by retaining those with the highest MS-DIAL total score. Detected synthetic drugs were removed from the dataset due to their lack of relevance to metabolite changes in response to the experimental treatment. Regarding all the features not positively identified (unknowns), following removal based on the previously mentioned sample max/blank average, unknowns that generated both m/z and MS2 data were retained and are reported separately. All sample peak heights (semi-quantitative) were normalized to the metabolite total ion chromatogram (mTIC) for each of the analyses performed (RPLC-Positive, RPLC -Negative).

To perform statistical analyses, processed peak intensity data above the 100,000 threshold for compound detection from the plant callus samples were uploaded onto MetaboAnalyst 6.0 (https://www.metaboanalyst.ca/). Peak intensity data was log transformed, and MetaboAnalyst was used to create PCA plots and heatmaps to describe unique metabolomic profiles between the sample groups. For metabolite expression and annotation (M2F, see below), a metabolite was considered present in each callus line if the median peak intensity across their replicates was ≥ 100,000.

### 2.4. Metabolite2Function (M2F) pipeline

#### 2.4.1. Searching PubMed Database

To identify literature that provides support for connecting metabolites to health functions, we created a pipeline to download, filter, and annotate relevant scientific paper title and abstract text data.

For each function, we created a set of closely related keywords to find paper matches on the PubMed database. Specifically, we searched using the phrase ({metabolite}[TIAB] AND ({keyword}[TIAB]) to find papers that had both the metabolite and the function of interest in either the title and/or abstract (Supplementary Table S1). Text data from titles and abstracts were downloaded from papers that matched this search. This search was repeated up to three times using metabolite synonyms fetched from PubChem (https://pubchem.ncbi.nlm.nih.gov/).

#### 2.4.2. Filtering Abstracts

After downloading the papers, extensive filtering on the abstracts is conducted to ensure correctly connecting the specifically searched metabolite to a function. Within each metabolite and its matching synonyms, duplicate papers across keywords for each function were removed.

Records were then filtered to remove non-matching metabolite entries. Specifically, abstracts with mismatched text referring to the target metabolite name or its synonyms were excluded. To achieve this, regular expression (regex) pattern matching was employed to identify and eliminate false positives that came from partial matches. For each metabolite, the regex pattern detected common mismatch scenarios, including hyphenated forms, enzyme-related extensions, amino acid conjugates, attached words, and “-yl” prefixes. By doing so, only abstracts containing exact and contextually relevant mentions of the metabolite were retained for subsequent analysis.

#### 2.4.3. GPT Metabolite Function Annotation

The extracted title and abstract text data were used as input for OpenAI text sentiment analysis using their GPT-4.1-mini model. For each paper, text data from each abstract was used as an input, and we used prompts tailored to each function to determine if the text provides information that the specific metabolite is relevant to that particular function (Supplementary Table S2). The general structure of the prompts is shown in Box 1. We limited the GPT response to Yes, No, or Unsure, and a short phrase less than 30 words describing its reasoning.

##### Box 1 – Anti-aging function prompt example

*Role*:

You are a rigorous scientist reviewing biological literature.

*Task*:

Evidence decision – decide whether the text below provides direct experimental evidence that the compound {metabolite} or {synonyms} has anti-aging properties.

Your answer must start with one of:

- Yes – direct evidence supports the claim
- No – direct evidence is missing or negative
- Unsure – cannot decide from the text Follow the decision with ONE short sentence (≤30 words) explaining whether the metabolite has anti-aging properties. State the metabolite name, the biomarker measured (e.g., collagen levels, MMP-1 inhibition), and the observed effect on aging markers.

*Clarifications*:

- “Direct” means the isolated compound was tested (in vitro / in vivo)
- If antiaging or anti-aging is not mentioned at all, answer “No”
- The metabolite or its synonyms must be verbatim

TEXT TO EVALUATE:

[*Paper title and abstract text*]

#### 2.2.4. Statistical Analysis

We performed statical analyses on annotation data using R version 4.2.3 (The R Foundation, r-project.org). For detailed analysis, a subset of 100 antioxidant and anti-senescence abstracts were used to test the performance of the GPT 4.1-mini model using the metrics of accuracy (overall correctness of the model) and precision (positive predictive value). These abstracts were annotated blindly by an individual human reviewer following the same evaluation prompt as above. Accuracy (1) and precision (2) were calculated as the following:

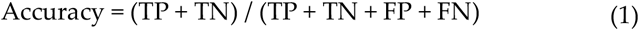

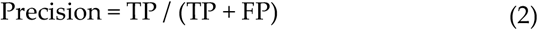

A true positive (TP) and true negative (TN) result occurs when the human annotator and GPT model agree, *i.e*., both human and GPT annotators label Yes or No, respectively, when evaluating the abstract text for function sentiment. A false positive (FP) result occurs when the GPT model labels the abstract as “Yes” and the human annotator labels as “No”. A false negative (FN) result occurs when the GPT model labels the abstract as “No” and the human annotator labels as “Yes”. We qualitatively compared the accuracy and precision for GPT and the individual human reviewer using a confusion matrix and bar plots. Subsequently, we will refer “Yes” labels as positive hits.

#### 2.4.5. Organism Classification

Following the metabolite function annotation, an organism-classification step was performed to restrict the dataset to studies conducted in humans or other mammals. For each positive hit, the title and abstract were submitted to a secondary GPT 4.1-mini prompt to categorize the paper. The model was required to assign the study to exactly one organism category (Humans, Mammals, Birds, Reptiles, Fish, Plants, Fungi, Bacteria, Viruses, Other) based on the provided text data. Positive hits that were not classified as Humans or Mammals were excluded from downstream analysis to ensure that retained annotations provided evidence relevant to human or mammalian biology.

### 2.5. 2,2-Diphenyl-1-picrylhydrazyl (DPPH) radical scavenging assay

Callus extracts were prepared as 10% (w/v) suspensions in water, homogenized, and filtered through 0.22 µm membranes. The radical scavenging activity of callus extracts and standard compounds (Vitamin C) were determined spectrophotometrically with the DPPH method [29]. The reaction was set by mixing 0.1 mL of tested samples with 0.9 mL of DPPH (81 mM in methanol) and stored in the dark at room temperature for 24 h. The absorbance was carried out by the BioTek Synergy LX spectrophotometer at 517 nm. Measurements were performed on the three biological triplicates. The results were expressed in micromolar equivalents of Vitamin C (L-ascorbic acid), which was calibrated using gradual concentrations from 0 to 500 mM.

### 2.6. Senescence-associated β-galactosidase (SA-β-gal)

Primary human epidermal melanocytes (HEMa; ATCC PCS-200-013) under oxidative stress conditions, were used to evaluate the anti-senescence effects of plant callus extracts (10% w/v in water). Cells cultured in Dermal Cell Basal Medium (ATCC PCS-200-030) were pretreated with callus extracts for 24 h, followed by exposure to hydrogen peroxide (H_2_O_2_, 200–300 µM) to induce stress-induced senescence. After 48–72 h, senescence-associated β-galactosidase (SA-β-gal) activity was assessed using the standard X-gal staining method at pH 6.0. Cells were fixed with 2% formaldehyde and 0.2% glutaraldehyde and incubated with the staining solution at 37 °C (no CO_2_) for 12–16 h. Blue-stained (SA-β-gal–positive) cells were counted under a light microscope, and the percentage of positive cells was compared between treated and control groups to deter-mine the anti-senes-cence efficacy of the callus extracts.

### 2.7. Anti-glycation

Plant extracts were prepared as 20% (w/v) suspensions in water, homogenized, and filtered through 0.22 µm membranes. Anti-glycation activity was tested using human serum albumin (HSA, 10 mg/mL) in PBS with glucose (0.22 M) as the glycating agent and sodium azide (0.1% w/v) to prevent microbial growth. Extracts were added at 10% (v/v) of the reaction volume. Controls lacking extract, glucose, or HSA were included. Reactions were incubated at 37 °C for 4 weeks, and advanced glycation end-products (AGE) formation was measured by fluorescence on a BioTek plate reader (Ex 360/40 nm, Em 460/40 nm). Activity was expressed as percent inhibition relative to glycated controls.

## 3. Results

### 3.1. Development of six plant cell cultures (calli)

We developed plant callus lines from five plants, namely flax (*Linum usitatissimum)*, hibiscus (*Hibiscus sabdariffa)*, shikakai (*Acacia concinna)*, tulsi (*Ocimum sanctum)*, and carrot (*Daucus carota*) (Figure 1a). The tobacco Bright Yellow 2 (BY-2) cell culture was used as a reference callus line [30]. These six plant calli represent six different taxonomic plant families and orders within the eudicot clade of flowering plants (Figure 1b). All calli have been developed from seedlings and were grown in the dark, except for carrot that was developed from taproot and was grown in a light/dark cycle. All calli were established cell cultures, i.e., 15-25 passages since their callus induction and reached stable morphological properties. Growth rates of the six calli ranged between 7%-11% per day (Figure 1c). Flax, tobacco, hibiscus, and shikakai were yellowish in color, whereas the tulsi and carrot calli were brownish and greenish, respectively (Figure 1a). Colored calli, i.e., tulsi and carrot, also showed a broader absorption of ultraviolet light than the yellowish callus cultures (Figure 1d).

**Figure 1.**
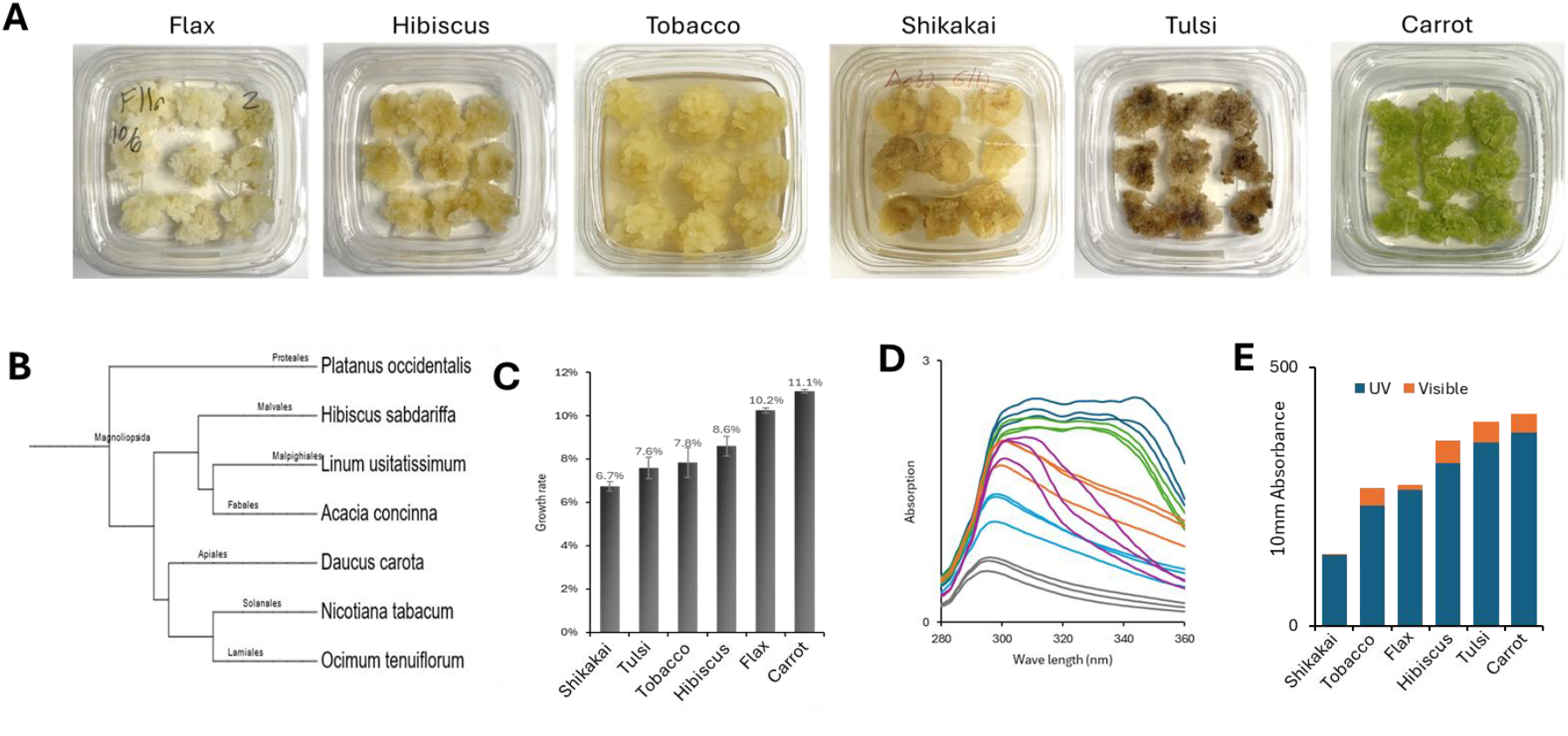
Development of plant calli. (**a**) Visual of the six plant callus cultures developed and investigated in this study. Calli are 28 days old since last passage. (**b**) Taxonomy tree of the six calli generated on phyloT v2 using NCBI Taxonomy. Platanus occidentalis (American sycamore) was used as an outgroup. The six plant calli belong to six different plant classes/orders (marked on the branches). (**c**) Growth rate of plant calli is shown as the percent of biomass increase per day. (**d**) UV absorbance spectrum of 10% callus extracts in 1:1 water:methanol solution. (**e**) Total light absorbance (UV + visible light) of plant calli water:methanol extracts.

### 3.2. Plant calli metabolomes clustered differently than their genomes

To compare the metabolomes of the six calli, we have subjected the calli to untargeted metabolomics using LC-MS/MS. Untargeted metabolomics have identified 6,902 unique mass features. 6,478 of these compounds passed the intensity cutoff in a minimum two biological replicates in at least one of the calli. Among the unique mass features, 181 have been identified as Metabolomics Standards Initiative level 1 or 2, i.e., MSI1 or MSI2, and 177 of these were present in at least one of the calli. Principal component analysis (PCA) separated the metabolomes by species, i.e., biological replicates of same species have been clustered together (Figure 2a). Hierarchical clustering further showed flax and hibiscus to be clustered closer to the tobacco reference line, shikakai to be more distant, and tulsi and carrot were grouped together and the most differentiated from the rest of the calli (Figure 2b). The metabolomics clustering of the plant calli was correlated with the clustering of the calli UV-absorbance spectrums, i.e., in both methods tulsi and carrot were clustered together and apart from the rest of the calli, while the flax and hibiscus calli clustered together with tobacco callus (Figures 2a-b versus Figures 2c-d).

**Figure 2.**
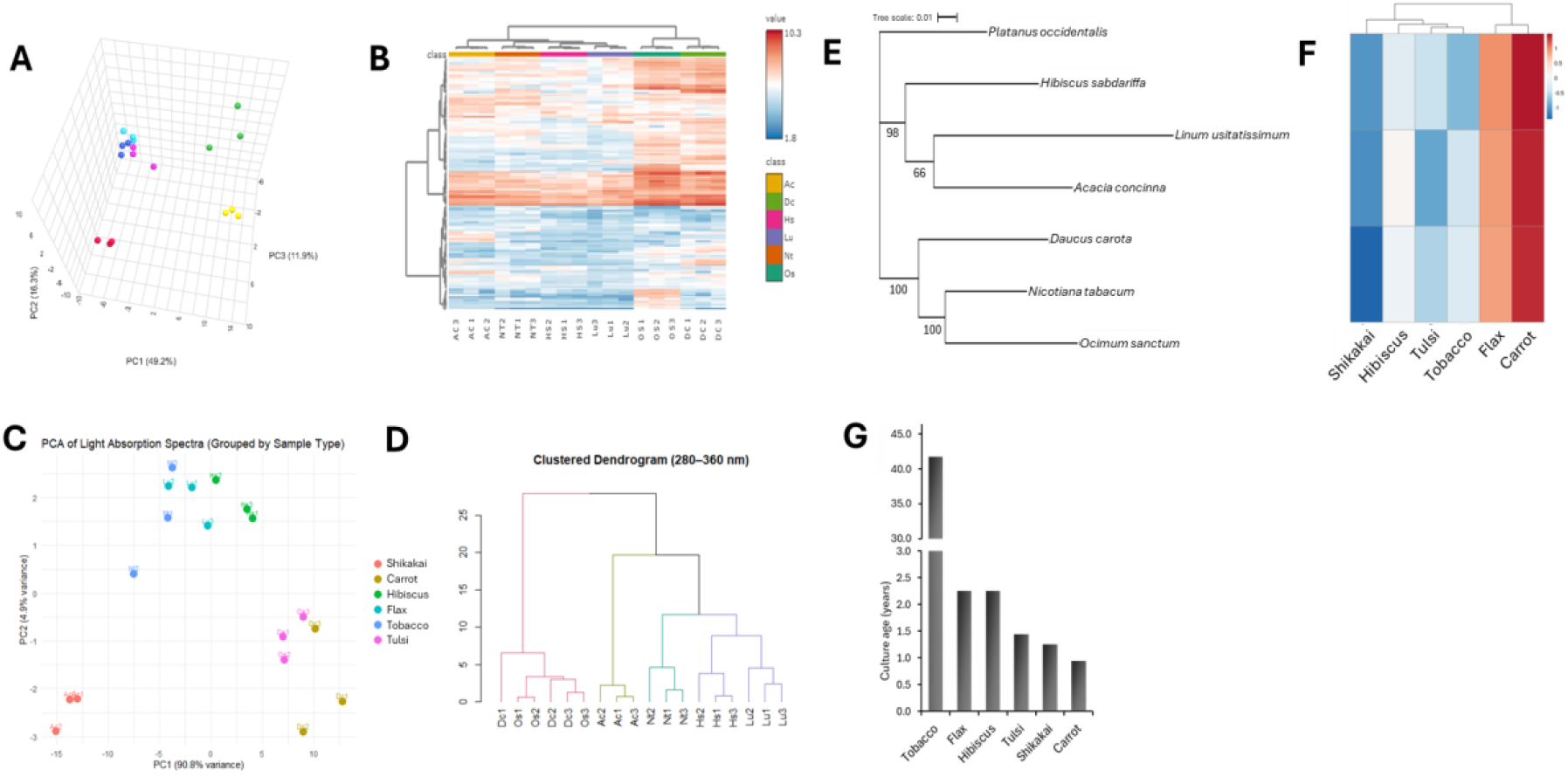
Plant callus metabolomes clustering. (**a**) PCA plot of the six plant callus metabolomes (three biological replicates per callus species). PCA plot is based on the LC-MS intensity of the 177 identified MSI1 and MSI2 metabolites. (**b**) Hierarchical clustering of the six plant callus metabolomes. PCA plot (**c**) and hierarchical clustering (**d**) of plant callus’ UV absorbance spectrums. (**e**) Evolutionary tree of the six plant calli based on their chloroplast genomic sequences, *Platanus occidentalis* was used as an outgroup. (**f**) Hierarchical clustering of the six plant calli based on growth rate presented in Figure 1c. (**g**) Callus age of the six plant calli.

Comparison of the metabolomic clustering to the taxonomy (Figure 1b) or to the genomic (Figure 2e) trees showed partial alignment. Like in the taxonomy and genomics trees, the pairs of flax/hibiscus and tulsi/carrot were found on separate clades in the metabolomics hierarchical clustering dendrogram (Figures 1b and 2e versus Figure 2b). However, while in the taxonomy/genomics phylogenetic trees the tobacco is grouped together with carrot and tulsi, in the metabolomics data it was grouped together with flax and hibiscus (Figures 2b and e). This result aligns with previous studies showing that metabolome-genome correlations exist but may vary across tissues, growth stages, and environmental conditions [31]. Plant calli can be considered as less differentiated cells, hence basic cell culture features, such as growth rate could play an important role in their metabolomes. To test this hypothesis, we quantified and compared the growth rate of the six plant calli to their metabolomes. Carrot and flax showed the fastest growth rates (10-11% per day), hibiscus, tobacco, and tulsi intermediate growth rate (8-9%), and shikakai had the slowest growth rate (∼7%) (Figure 1c). These growth rates do not correspond with the clustering patterns observed in the calli’s metabolomes (Figures 2b and f), suggesting that cellular growth rate is not a primary determinant of metabolomic variation across species.

Another biological feature that could affect plant callus metabolomes is the age of the calli, i.e., the time passed since the induction of the callus culture from the mother plant, which is indicative to the number of cell divisions and the differentiation potential of the callus (the ability to develop into other cell types). The age of the carrot callus was about one year old, shikakai and tulsi were about 1.5 years old, and flax and hibiscus were a bit over 2 years (Figure 2g). In comparison to these cultures, the tobacco BY-2 culture was developed about 4 decades ago (Figure 2g). Comparing the callus age to the metabolomic data showed some correlation but not a complete one (Figures 2b and 2g). For example, flax and hibiscus have similar age and show the closest metabolomic profiles (Figures 2b and g). Additionally, among our developed calli, flax and hibiscus are the oldest ones, and their metabolomes were clustered together with the 4-decade old tobacco line. Albeit these correlations, tulsi and carrot which their metabolomes clustered together and separately from the rest of the calli, showed a similar age to that of the shikakai. Additionally, if callus age was a determining factor of calli metabolomes derived from different species, one might have expected to find the tobacco BY-2’s metabolome to be distinguished from the rest of the calli, as the number of its cell divisions substantially supersedes that of the other calli. Overall, these results suggest that neither genetics, callus age, nor callus growth rate, can be used individually to classify the plant calli metabolomes.

### 3.3 Metabolite enrichment levels in the six plant calli

Among the six calli, tulsi and carrot were significantly enriched in metabolites, i.e., having the highest number of identified metabolites and the highest level of metabolite intensity (Figures 3a-d). In comparison, flax and hibiscus were relatively depleted of metabolites, both in the number and intensity of metabolites (Figures 3a-d). Shikakai and tobacco calli were found to be in the middle in terms of the number and intensity level of their metabolites (Figures 3a-d). Metabolite enrichment level was not correlated with moisture content of the samples (Figure 3e), suggesting the proportion of dry matter is not the main reason for the metabolite content in the different calli.

**Figure 3.**
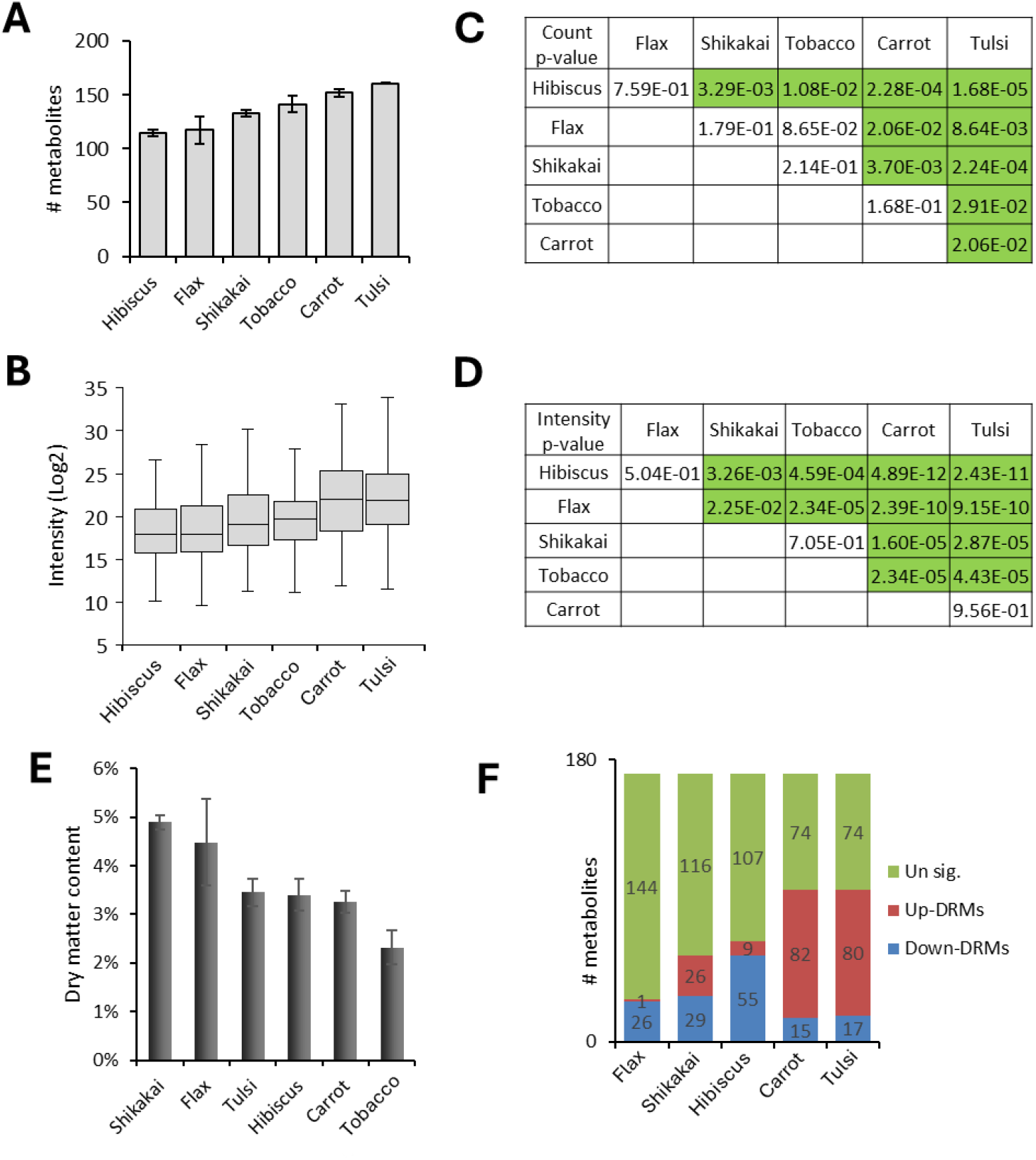
Metabolite enrichment levels in the six plant calli. (a) Average number of MSI1 and MSI2 metabolites identified in each of the plant calli. (b) Boxplot of LC-MS intensity of MSI1 and MSI2 metabolites in the six plant calli. C-D. Student’s t-TEST between the indicated plant calli’s number of identified metabolites (c) or intensity level (d). Green boxes represent significant difference (p < .05). (e) Dry matter content in plant calli. (f) Stacked-column graph showing the number of upregulated-DMRs, downregulated-DMRs, and unregulated-DMRs in the indicated calli versus the tobacco callus.

These findings were correlated with the number of differentially regulated metabolites (DRM) that were significantly upregulated (up-DRMs) or downregulated (down-DRMs) in the different calli versus the tobacco reference line (Figure 3f). In comparison to the tobacco callus, carrot and tulsi calli were relatively enriched in up-DRMs, hibiscus and flax were relatively enriched in down-DRMs, and shikakai had about the same number of up- and down-DRMs (Figure 3f). Accordingly, the metabolomes of our recently developed callus lines were found to be relatively enriched or depleted in comparison to the established tobacco four-decades-old callus line, further supporting that callus age is not necessarily playing a substantial factor in the metabolome of plant calli derived from different species.

### 3.4. Metabolite differentiation levels in the six plant calli

To learn about the level of metabolite consistency or differentiation between plant calli we next identified DRMs amongst the six plant calli. We then plotted the metabolites on a ‘Robustness’ scale that ranges between 0-15; zero Robustness refers to metabolites that were not differentiated at all between any of the calli, whereas a Robustness score of 15 refers to metabolites that were differentiated between all six calli (i.e., 15 pairwise comparisons). A distribution plot of metabolites over the Robustness score showed that there are no metabolites with a zero or one Robustness scores, i.e., there are no metabolites that were 100% consistent among the six calli (Figure 4a). Second, it shows that most metabolites (63%) have a Robustness score higher than 7, i.e., most metabolites are differentiated between at least eight pairwise callus comparisons (Figure 4a). Additionally, we found that 70% of metabolites showed a different scale of metabolite intensity in at least four calli out of the six of them (Figure 4b). Metabolic pathway analysis found metabolites involved in amino acids metabolism to be significantly enriched among mildly-differentiated metabolites (Robustness score 2-6), whereas highly-differentiated metabolites (Robustness score 10-12) were enriched in starch and sucrose metabolism (Figures 4c-d). To further explore the differentiation of metabolite abundance among the six plant calli, we plotted the metabolites according to their occurrence frequency in the plant calli, i.e., ‘Prevalence’ scale. A Prevalence of one and six refer to metabolites that were specifically detected in a single plant callus or ubiquitously found in all plant calli, respectively. Using the Prevalence scoring system we found that 55% of metabolites are ubiquitous (have been produced within all six calli), 35% of metabolites are common (have been produced in 2-5 calli), and 10% of metabolites are unique (produced in a single callus species) (Figure 4e). Additionally, we found that the metabolite intensity level of unique metabolites, i.e., detected in a single callus, to be relatively higher than most commonly produced metabolites (Figure 4f). Overall, these findings suggest that most metabolites are highly differentiated in their abundance among the six plant calli, and that unique metabolites may contribute to a significant portion of the plant calli metabolome.

**Figure 4.**
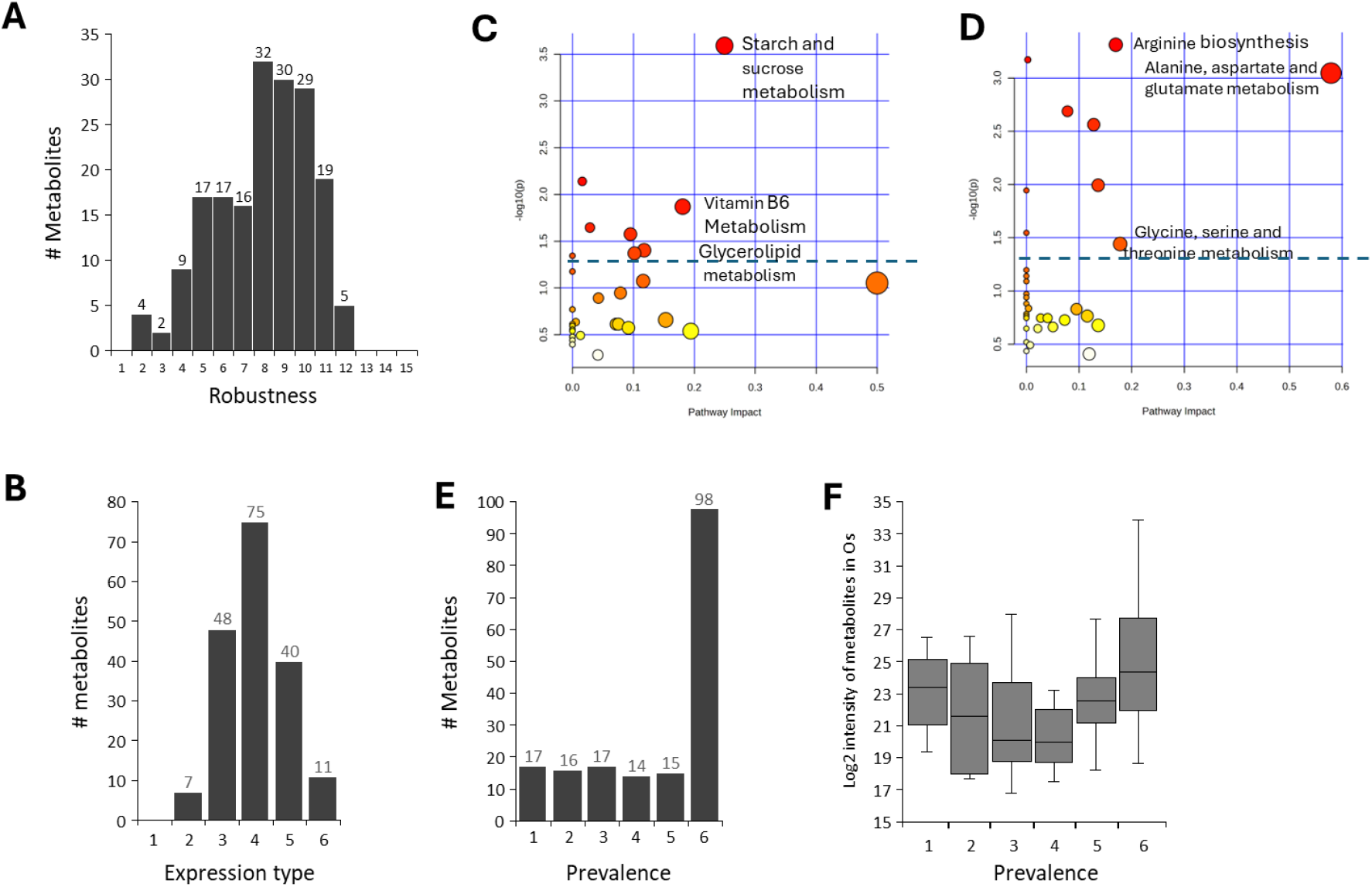
Metabolites differentiation level in the six plant calli. (a) Distribution plot of the number of metabolites over Robustness level. Robustness score of 1 and 15 represent metabolites that were differentially expressed in one or 15 callus pairwise comparisons. (b) Distribution plot of the number of metabolites found at each expression type. Expression type 1 and 6 represent metabolites that were differentially produced in neither or all the six calli, respectively. Heat scatter plots of the representation of metabolic pathways among metabolites that relatively highly (c) or lowly (d) differentiated among the six plant calli, i.e., having a Robustness score bigger than 9 or smaller than 7, respectively. Size and color of the dots represent the number of metabolites and the p-value in each metabolic pathway, respectively. (e) Distribution plot of the number of metabolites over the Prevalence scale (metabolite occurrence rate); score of 1 and 6 refer to metabolites found in a single callus or in all six calli, respectively. (f) Boxplot of metabolite LC-MS intensity in the tulsi callus over Prevalence scale.

### 3.5. Annotating metabolites to biological functions beneficial to human health

One of our central goals is determine whether the metabolites present in our plant calli have beneficial properties in supporting human health. Because metabolite ontology databases are currently missing or insufficient, we set out to develop a metabolite ontology pipeline. Our custom metabolite annotation pipeline, Metabolite2Function (M2F), combines large-scale PubMed searches with OpenAI’s GPT LLM to annotate paper title and abstract text data (Figure 5a). Only the title and abstract were included in the pipeline, rather than full manuscript, to be relied on freely accessible and highly relevant data, as well as to enable faster and more cost-efficient annotation. This design depends on availability, relevance, and efficiency, and it facilitates adapting the pipeline for other metabolites or functions of interest.

**Figure 5.**
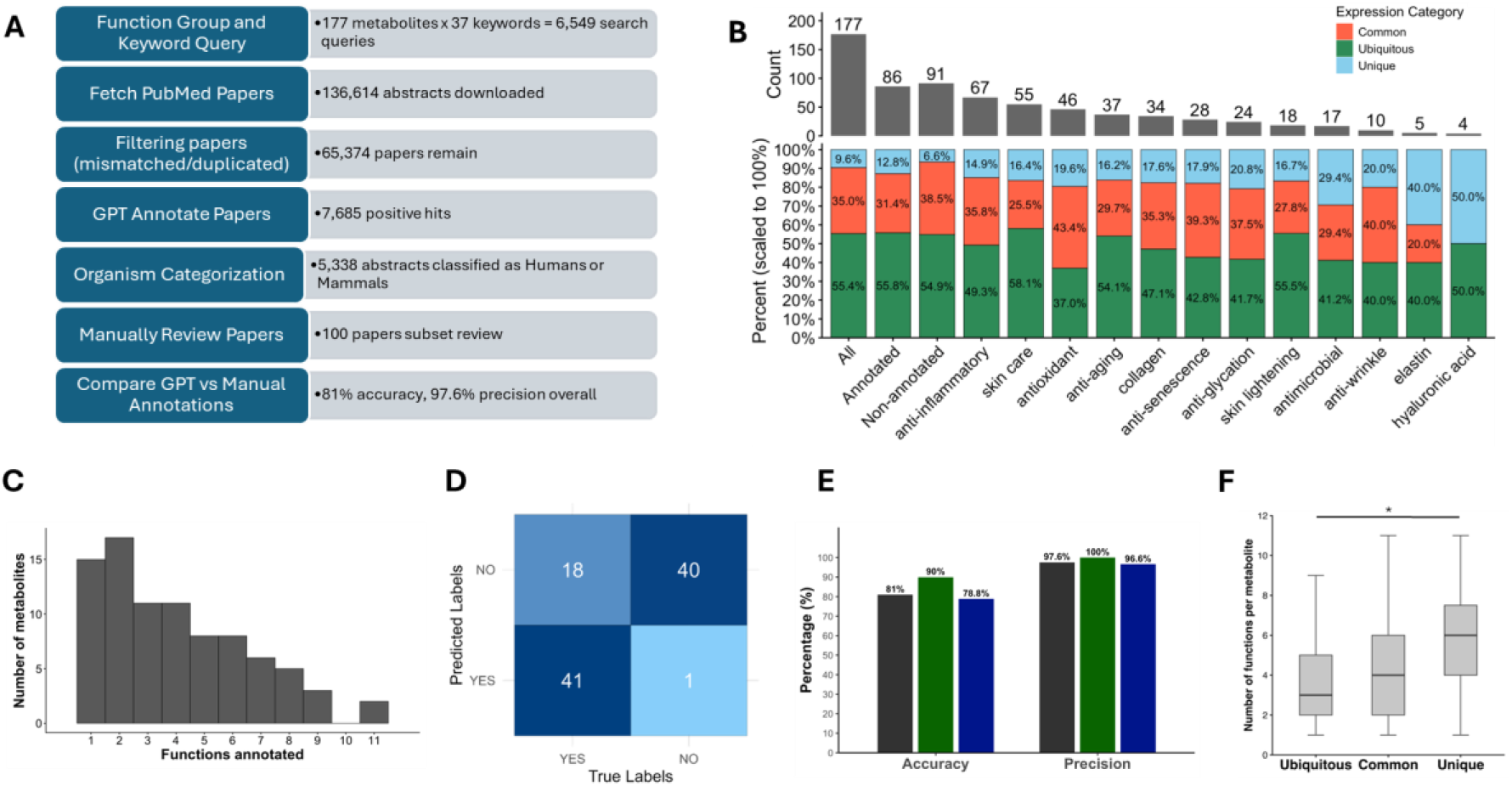
Annotating metabolites to desired biological functions. (**a**) Flowchart of the annotation pipeline with relevant numbers. (**b**) Enrichment of bioactive metabolites among callus-specific metabolites. Stacked column graphs showing the number of unique (blue), common (orange), and ubiquitous (green) metabolites according to the Prevalence score among all metabolites and different functional annotation groups. (**c**) Histogram of functions annotated per metabolite. (**d**) The confusion matrix represents the congruence between human annotations, i.e., manually reviewed (True labels) and the GPT 4.1-mini model (Predicted labels). (**e**) Bar graphs depicting accuracy and precision of model annotations of total (black), anti-senescence (dark green), and antioxidant (dark blue) annotated functions. (**f**) Box plots showing the distribution of the number of annotated functions per metabolite for ubiquitous, common, and unique metabolites (^*^ Tukey’s HSD p = .015).

Untargeted metabolomic analysis on the plant callus samples identified 177 MSI 1 and MSI 2 metabolites. These MSI 1 and MSI 2 metabolites were searched against the PubMed database to discover abstracts that support whether they have beneficial properties important in skin health. We were specifically interested in the following eleven functions that promote anti-aging, anti-glycation, anti-inflammation, antimicrobial, antioxidant, anti-senescence (cellular), anti-wrinkle, collagen, elastin, hyaluronic acid, skin care, and skin lightening.

We used function-specific prompts to identify papers that provided support that a given metabolite contributed to health benefits. For the eleven chosen biological functions, we used 37 keywords and 177 MSI 1 and MSI 2 metabolites found in our dataset resulting in 6,549 PubMed search queries. We limited our search results to the top 100 most relevant papers per function, resulting in 136,614 papers found (Figure 5a). After filtering duplicates within function groups and non-perfect matched metabolites we ended up with 65,374 papers to annotate. GPT model 4.1-mini identified 7,685 positive hits that provided evidence in linking metabolites to specific functional annotations. Of these positive hits, 5,338 abstracts were classified as relevant to Humans or mammals. Overall, we have annotated 87 beneficial (bioactive) metabolites in our six callus samples, which is 49% of the metabolites in the MSI1 and MSI2 groups (Figure 5b). The median number of positive hits and functions per metabolite were 7 and 3, respectively. Some metabolites, such as (-)-Epigallo-catechin-3-gallate, had 11 annotated functions and 200 positive hit papers (Figure 5c).

The results from comparing GPT and individual human annotation of the anti-senescence and antioxidant 100 paper subset is shown in Figure 5d. We calculated an overall accuracy of 81% and an overall precision of 97.6%. For our anti-senescence and antioxidant subset, we calculated an accuracy of 90% and 78.8%, and a precision of 100% and 96.6%, respectively (Figure 5e). These results indicate that the model is performing accurately and most of the positively annotated papers by the GPT model were classified correctly. The model has a lower accuracy determining true negatives; however, a high precision for true positives indicates that papers annotated as such are reliable.

### 3.6. Bioactive metabolites are enriched among callus-specific metabolites

The plant calli with the highest and lowest beneficial (bioactive) metabolites were tulsi with 76 metabolites and flax with 43 metabolites, respectively. The proportion of unique metabolites among beneficially annotated metabolites was significantly higher than that of unannotated metabolites, 12.8% versus 6.6%, respectively (Figure 5b). In particular functional annotations, the proportion of unique metabolites was even higher, such as 19.6% and 17.6% in antioxidant and pro-collagen, respectively (Figure 5b). Additionally, unique metabolites had the highest number of annotated functions per metabolite (median = 6), compared with common metabolites (median = 4), and robust metabolites (median = 3) (Figure 5f). An analysis of variance showed that Prevalence score had a significant effect on the number of annotated functions per metabolite, *F*(2,84) = 4.51, *p* = 0.0137. Post hoc analyses using Tukey’s HSD test indicate that unique metabolites were associated with a significantly higher number of annotated functions than robust metabolites (mean difference = 2.35, 95% CI [0.38, 4 .32], *p* = 0.015), whereas common metabolites did not differ significantly from either group. Together, these results demonstrate the significant portion of beneficial bioactive metabolites among species-specific callus metabolomes.

### 3.7. Spermidine function case study

To evaluate how well the text-mining and GPT annotation steps captured function-relevant annotations, we focused on analyzing papers for the metabolite spermidine. Spermidine is natural polyamine that is critical to the maintenance of cellular homeostasis and has been seen to extend lifespan and health span within yeast, nematodes, flies, and mice [32]. It is also known to have anti-inflammatory and antioxidant properties [33,34] so we wanted to investigate if our pipeline can map this metabolite to any of our a priori functions. Across PubMed queries combining “spermidine” along with function-specific keywords, we downloaded 2,327 papers. After paper duplicate removal and regex-based filtration, 1,619 abstracts remained (Figure 6a). Applying our prompt using GPT 4.1-mini model yielded 161positive hits, corresponding to an estimate of 9.9% among filtered abstracts. Of these positives, 137 or 85% of abstracts were classified as Humans/Mammals based on the organism-context prompt.

**Figure 6.**
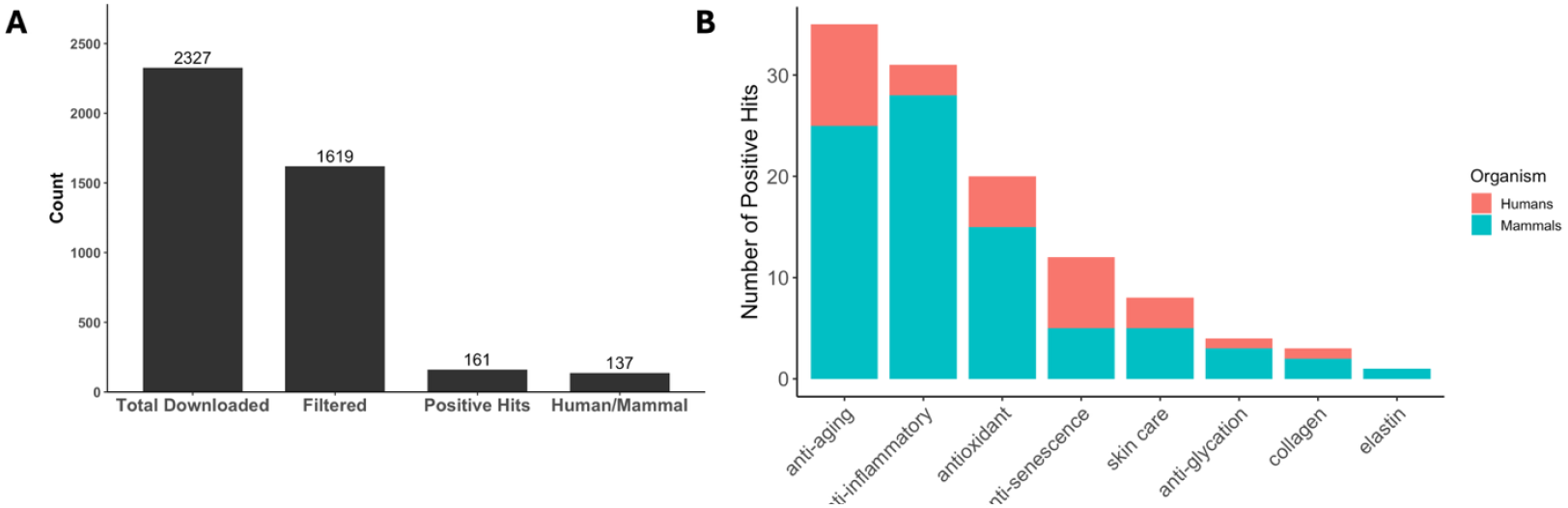
Spermidine metabolite paper count and distribution of positive hits. (**a**) Count of total spermidine papers downloaded from PubMed matching function keywords, number papers remaining after filtering, number of positive hit papers, and how many of these were related to Human or mammal studies. (**b**) Number of positive hit papers by function, separated by studies focusing on Humans or Mammals.

The distribution of positive hits by function (Figure 6b) shows that spermidine is most frequently supported for anti-aging and anti-inflammatory roles, with additional functions in antioxidant, anti-senescence, anti-glycation, skin care, collagen, and only a few positives mapped to elastin. Notably, the human and mammal subset of papers contribute to a substantial portion of the positives, suggesting that the literature support is not restricted to plant or invertebrate models. The spermidine profile is useful as an internal check for our pipeline as it recapitulates expected biology with stronger signals in anti-aging, anti-inflammatory, and antioxidant functions. This lends confidence that our pipeline retrieves relevant evidence allowing us to connect our plant callus metabolites to biological functions.

### 3.8. The abundance of antioxidants and anti-senescence metabolites in plant calli correlate with their corresponding biochemical activities

Next, we wanted to check if the LC-MS intensity (i.e., abundance) of annotated metabolites correlates with their correspondent biochemical or biological activities. For example, does the metabolite intensity of the overall metabolites annotated to antioxidation (i.e., antioxidants) in the different calli correlate with the antioxidant capacity of the calli. To that end, we first measured the antioxidant activity of callus extracts derived from the six calli using the DPPH method. Then, we compared antioxidant activity to the level of LC-MS intensities of metabolites annotated as antioxidants. This analysis found a positive and significant correlation between metabolite intensity of annotated antioxidants and the antioxidant activity among the six calli, Pearson’s r = 0.88, *p* < .0206 (Figure 7a).

**Figure 7.**
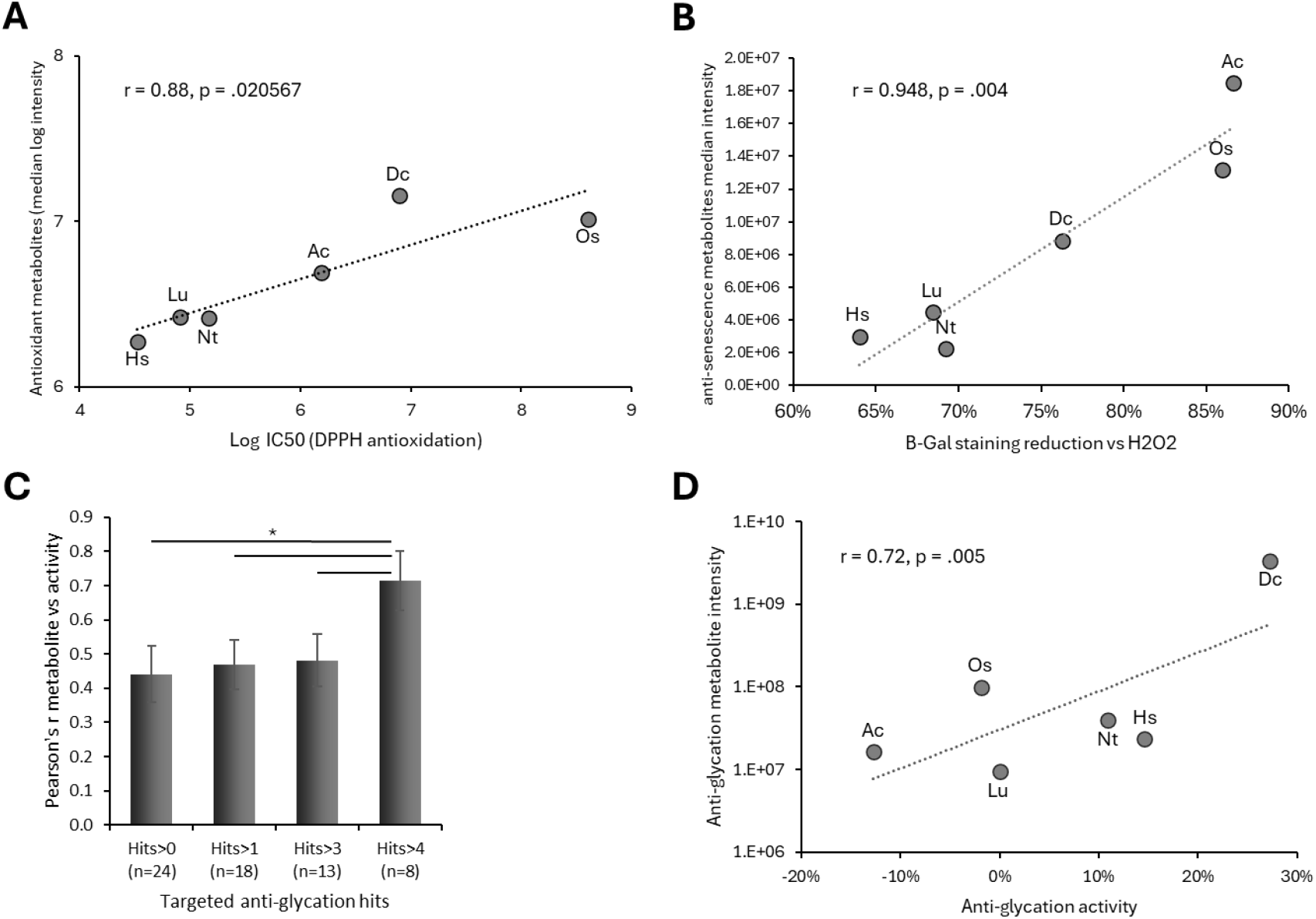
Metabolite annotation scorings correlate with in vitro activity. (**a**) Scatter plot of the median intensity level of all annotated antioxidants in the six plant calli versus the antioxidant capacity measure by DPPH in the six calli. (**b**) Scatter plot of the median intensity level of all annotated anti-senescence metabolites in the six plant calli versus the anti-senescence activity of the plant calli on human melanocytes. Anti-senescence activity was measured by the suppression level of senescence, quantified by beta-gal staining, induced by hydrogen peroxide. r represents Pearson’s correlation coefficient. (c) Pearson’s correlation coefficient values of total intensity of anti-glycation annotated metabolites over the anti-glycation activity of human serum albumin by the plant calli. Correlation coefficient was measured for metabolites with the indicated minimum number of positive anti-glycation annotation hits (^*^, p-value < 0.006; n = number of metabolites in the category). Ac, Dc, Hs, Lu, Nt, and Os refer to shikakai, carrot, hibiscus, flax, tobacco, and tulsi, respectively. (**d**) Scatter plot of metabolite intensity level of annotated anti-glycation metabolites (>4 positive hits) over anti-glycation activity of human serum albumin by plant callus extracts.

Antioxidation is a robust biochemical activity of many metabolites. Hence, the overall enrichment level of metabolites in the different calli could also infer on the antioxidant capacity of the samples. Next, we wanted to compare the intensity of metabolites annotated to a biological function with less annotated metabolites and thus less influenced by the overall metabolome intensity of the samples. To that end, we chose the anti-senescence function, which includes 10 annotated metabolites among either of our plant calli. Anti-senescence effect of the plant calli on human cells was measured by subjecting callus extracts on human melanocytes induced for senescence by hydrogen peroxide. Cellular senescence was monitored with beta-galactosidase. All six calli showed anti-senescence effect where its strength was correlated with the metabolite intensity of the annotated anti-senescence metabolites in the corresponding samples (Figure 7b). First, a strong and significant correlation was found between anti-senescence metabolite intensity and anti-senescence activity in the six calli (*p* = 0.004). Second, unlike with antioxidants, in this case, the callus with the most anti-senescence metabolites and activity was the shikakai, despite having a moderate overall metabolome enrichment level among the six calli. These findings suggest that the intensity of metabolites annotated by our novel model can infer on the biological activity of the profiled biological samples (e.g., plant calli).

### 3.9. Anti-Glycation Activity Correlates with Stringency-Filtered Metabolomics Rankings

We next compared metabolomics-based rankings with anti-glycation activity. Protein glycation is a harmful process that leads to the formation of advanced glycation end-products (AGEs), which accelerate aging and contribute to wrinkles, tissue stiffness, and chronic disease [35]. The anti-glycation activity of plant calli was assessed by measuring inhibition of human serum albumin glycation. Among the six calli tested, three exhibited measurable anti-glycation activity, with the carrot callus showing the strongest effect, reaching 27% inhibition of glycation (Figure 7c).

We identified 24 metabolites with at least one literature-supported association (i.e., positive hits) with anti-glycation activity (Figure 5b). When comparing the total intensity of all annotated anti-glycation metabolites in the plant calli with their measured anti-glycation activity, a mild correlation was observed (Figure 7c). However, increasing the stringency of the annotation threshold improved the correlation: requiring a minimum of four positive citation hits yielded a moderate correlation coefficient (Pearson’s r = 0.715, p < .005) between the summed metabolite intensities and the in vitro anti-glycation activity (Figures 7c-d), indicating that filtering metabolites by stronger literature support improves the ability of metabolomics data to predict functional bioactivity.

## 4. Discussion

This study advances our understanding of the comparative metabolomics of plant calli and highlights their potential as rich sources of bioactive compounds with significant health-related applications. By profiling the metabolomes of six distinct plant calli—*Acacia concinna, Daucus carota, Hibiscus sabdariffa, Linum usitatissimum, Ocimum sanctum*, and *Nicotiana tabacum* (BY-2)—we identified a diverse array of metabolites with unique abundance patterns and characteristics. Our findings demonstrate that metabolomic profiles are shaped by a complex interplay of factors and do not strictly reflect genetic relationships, culture age, or growth rate.

Notably, most metabolites varied substantially in abundance among calli, indicating pronounced metabolic specialization. Metabolites unique to individual calli were present at relatively higher abundance and were disproportionately enriched in functionally annotated bioactivities. Moreover, unique metabolites exhibited a greater number of annotated functions per metabolite compared with common or ubiquitous metabolites. Collectively, these observations underscore the importance of metabolomic variability among plant calli and highlight species-specific callus metabolomes as concentrated sources of functionally enriched bioactive compounds with significant potential for targeted discovery and application [20,21].

A key innovation of this study is the development of Metabolite2Function (M2F), a novel and efficient metabolite annotation pipeline that leverages large language model (LLM)–based AI to assign putative biological functions to metabolites using abstract-level text mining. While AI-driven approaches are increasingly being adopted to support large-scale literature screening and systematic reviews [23,24,36], M2F extends this paradigm directly into functional metabolomics. By enabling automated, scalable annotation of metabolite–function relationships, M2F bridges the gap between metabolite identification and biological interpretation. This framework allowed us to map metabolite profiles to key biological processes and functional categories, revealing compelling evidence that plant calli represent a valuable source of anti-aging and other health-promoting compounds. The observed relationships between metabolite abundance and biological relevance further demonstrate the predictive power of metabolomic profiling when coupled with intelligent annotation tools such as M2F.

## 5. Conclusions

Overall, this research emphasizes both the biotechnological value of plant tissue cultures as sustainable and consistent platforms for bioactive compound production, and the critical role of AI-enabled pipelines such as M2F in unlocking functional insights from metabolomics data. Together, these advances pave the way for future focused studies aimed at experimentally validating the functional activities of identified metabolites, refining culture conditions for targeted metabolite synthesis, and expanding the application of plant-derived compounds in pharmaceuticals, nutraceuticals, and cosmetics. Furthermore, the integration of systematic metabolite-to-function mapping enables the rational selection of plant cultures to design personalized formulations tailored to specific biological effects, such as antioxidant, anti-inflammatory, or skin-beneficial activities. Continued investigation into plant calli, supported by integrative tools like M2F, holds significant promise for the discovery of novel and sustainable solutions to promote human health and well-being.

## Abbreviations

The following abbreviations are used in this manuscript:

LLM: Large language models
M2F: Metabolite2Function
SA-β-: 
Gal: Senescence-associated β-galactosidase
HEMa: Human epidermal melanocytes
AGE: Advanced glycation end-products
ATCC: American Type Culture Collection
TP: True positive
TN: True negative
FP: False positive
FN: False negative
HSA: Human serum albumin
PCA: Principal component analysis
DRM: Differentially regulated metabolite
Up-: 
DRMs: Significantly upregulated DRMs
Down-: 
DRMs: Significantly downregulated DRMs

## Author Contributions

Conceptualization, A.Z. and N.N.; methodology, M.R.P., A.Z., B.S.L., L.L.D., E.S.R., D.S., and D.T.A.; software, M.P.; formal analysis, A.Z., and M.R.P.; investigation, M.R.P., A.Z., B.S.L., L.L.D., D.S., and M.P.; data curation, M.R.P. and A.Z.; writing—original draft preparation, A.Z. and M.R.P.; writing— review and editing, A.Z., M.R.P., E.S.R., N.N., and D.T.A.; supervision, A.Z., and N.N.; funding acquisition, N.N. All authors have read and agreed to the published version of the manuscript.

## Institutional Review Board Statement

Not applicable.

## Informed Consent Statement

Primary Epidermal Melanocytes; Normal, Human, Adult (HEMa) (ATCC PCS-200-013) were purchased from American Type Culture Collection (ATCC). The cells were obtained from anonymized human donors with informed consent by the supplier. According to institutional guidelines, research involving commercially available de-identified primary human cells does not constitute human subjects research and did not require ethical review or approval.

## Data Availability Statement

The original contributions presented in this study are included in the article. Further inquiries can be directed to the corresponding authors.

## Acknowledgments

The authors would like to thank May Bar-Yosef and Marcelo Rotsztejn for their assistance with procurement and administrative support. The authors have reviewed and edited the output and take full responsibility for the content of this publication.

## References

1. Shen S, Zhan C, Yang C, Fernie AR, Luo J. Metabolomics-centered mining of plant metabolic diversity and function: Past decade and future perspectives. Molecular Plant. 2023. doi:10.1016/j.molp.2022.09.007

2. Chen JT. Phytochemical omics in medicinal plants. Biomolecules. 2020. doi:10.3390/biom10060936

3. Veiga M, Costa EM, Silva S, Pintado M. Impact of plant extracts upon human health: A review. Critical Reviews in Food Science and Nutrition. 2020. doi:10.1080/10408398.2018.1540969

4. Hossain MS, Wazed MA, Asha S, Amin MR, Shimul IM. Dietary Phytochemicals in Health and Disease: Mechanisms, Clinical Evidence, and Applications—A Comprehensive Review. Food Science and Nutrition. 2025. doi:10.1002/fsn3.70101

5. Dillard CJ, Bruce German J. Phytochemicals: Nutraceuticals and human health. Journal of the Science of Food and Agriculture. 2000. doi:10.1002/1097-0010(20000915)80:12<1744::AID-JSFA725>3.0.CO;2-W

6. Matsuda F, Hirai MY, Sasaki E, Akiyama K, Yonekura-Sakakibara K, Provart NJ, et al. AtMetExpress development: A phytochemical atlas of Arabidopsis development. Plant Physiol. 2010;152(2). doi:10.1104/pp.109.148031

7. Schneider GF, Salazar D, Hildreth SB, Helm RF, Whitehead SR. Comparative Metabolomics of Fruits and Leaves in a Hyperdiverse Lineage Suggests Fruits Are a Key Incubator of Phytochemical Diversification. Front Plant Sci. 2021;12. doi:10.3389/fpls.2021.693739

8. Namdar D, Mazuz M, Ion A, Koltai H. Variation in the compositions of cannabinoid and terpenoids in Cannabis sativa derived from inflorescence position along the stem and extraction methods. Ind Crops Prod. 2018;113. doi:10.1016/j.indcrop.2018.01.060

9. Liu C, Wang L, Wang J, Wu B, Liu W, Fan P, et al. Resveratrols in Vitis berry skins and leaves: Their extraction and analysis by HPLC. Food Chem. 2013;136(2). doi:10.1016/j.foodchem.2012.08.017

10. Lin YL, Juan IM, Chen YL, Liang YC, Lin JK. Composition of polyphenols in fresh tea leaves and associations of their oxygen-radical-absorbing capacity with antiproliferative actions in fibroblast cells. J Agric Food Chem. 1996;44(6). doi:10.1021/jf950652k

11. Krivokapić S, Pejatović T, Perović S. Chemical characterization, nutritional benefits and some processed products from carrot (Daucus carota l.). Agriculture and Forestry. 2020;66(2). doi:10.17707/AgricultForest.66.2.18

12. Kocaadam B, Şanlier N. Curcumin, an active component of turmeric (Curcuma longa), and its effects on health. Crit Rev Food Sci Nutr. 2017;57(13). doi:10.1080/10408398.2015.1077195

13. Wijerathna-Yapa A, Hiti-Bandaralage J, Pathirana R. Harnessing metabolites from plant cell tissue and organ culture for sustainable biotechnology. Plant Cell, Tissue and Organ Culture. 2025. doi:10.1007/s11240-025-03180-6

14. Verma P, Khan SA, Alhandhali AJA, Parasharami VA. Bioreactor upscaling of different tissue of medicinal herbs for extraction of active phytomolecules: A step towards industrialization and enhanced production of phytochemicals. In: Plant Growth Regulators: Signalling under Stress Conditions. 2021. doi:10.1007/978-3-030-61153-8_21

15. Gautheret RJ. Plant tissue culture. Endeavour. 1948;7(26). doi:10.1385/0-89603-127-6:499

16. Verdeil JL, Alemanno L, Niemenak N, Tranbarger TJ. Pluripotent versus totipotent plant stem cells: dependence versus autonomy? Trends Plant Sci. 2007;12(6). doi:10.1016/j.tplants.2007.04.002

17. Weigel D, Jürgens G. Stem cells that make stems. Nature. 2002. doi:10.1038/415751a

18. Jiang F, Feng Z, Liu H, Zhu J. Involvement of plant stem cells or stem cell-like cells in dedifferentiation. Frontiers in Plant Science. 2015. doi:10.3389/fpls.2015.01028

19. Torres KC. Overview of Callus (Tissue) and Organ Culture. In: Tissue Culture Techniques for Horticultural Crops. 1989. doi:10.1007/978-1-4615-9756-8_5

20. Kim JS, Sato M, Kojima M, Asrori MI, Uehara-Yamaguchi Y, Takebayashi Y, et al. Multi-omics signatures of diverse plant callus cultures. Plant Biotechnology. 2024;41(3). doi:10.5511/plantbiotechnology.24.0719a

21. Szűcs Z, Cziáky Z, Volánszki L, Máthé C, Vasas G, Gonda S. Production of Polyphenolic Natural Products by Bract-Derived Tissue Cultures of Three Medicinal Tilia spp.: A Comparative Untargeted Metabolomics Study. Plants. 2024;13(10). doi:10.3390/plants13101288

22. Gilardi F, Alizadeh M, Kubli M. ChatGPT outperforms crowd workers for text-annotation tasks. Proc Natl Acad Sci U S A. 2023;120(30). doi:10.1073/pnas.2305016120

23. Issaiy M, Ghanaati H, Kolahi S, Shakiba M, Jalali AH, Zarei D, et al. Methodological insights into ChatGPT’ screening performance in systematic reviews. BMC Med Res Methodol. 2024;24(1). doi:10.1186/s12874-024-02203-8

24. Xu S, Zhao Z, Liu X, Meng XL. A comparative study of screening performance between abstrackr and GPT models: Systematic review and contextual analysis. BMC Medical Informatics and Decision Making. Bio-Med Central Ltd; 2025. doi:10.1186/s12911-025-03138-w

25. Oliveira SMR, Girol AP, Nissapatorn V, Pereira M de L. Bioactive Compounds Derived from Plants and Their Medicinal Potential. Pharmaceuticals. 2025. doi:10.3390/ph18111732

26. Bonini P, Kind T, Tsugawa H, Barupal DK, Fiehn O. Retip: Retention Time Prediction for Compound Annotation in Untargeted Metabolomics. Anal Chem. 2020;92(11). doi:10.1021/acs.analchem.9b05765

27. Tsugawa H, Cajka T, Kind T, Ma Y, Higgins B, Ikeda K, et al. MS-DIAL: Data-independent MS/MS deconvolution for comprehensive metabolome analysis. Nat Methods. 2015;12(6). doi:10.1038/nmeth.3393

28. Sumner LW, Amberg A, Barrett D, Beale MH, Beger R, Daykin CA, et al. Proposed minimum reporting standards for chemical analysis: Chemical Analysis Working Group (CAWG) Metabolomics Standards Initiative (MSI). Metabolomics. 2007;3(3). doi:10.1007/s11306-007-0082-2

29. Blois MS. Antioxidant determinations by the use of a stable free radical [10]. Nature. 1958. doi:10.1038/1811199a0

30. Katō K, Shiozawa Y, Yamada A, Nishida K, Noguchi M. A Jar Fermentor Culture of Nicotiana tabacum L. Cell Suspensions. Agric Biol Chem. 1972;36(6). doi:10.1271/bbb1961.36.899

31. Chan EKF, Rowe HC, Hansen BG, Kliebenstein DJ. The complex genetic architecture of the metabolome. PLoS Genet. 2010;6(11). doi:10.1371/journal.pgen.1001198

32. Madeo F, Bauer MA, Carmona-Gutierrez D, Kroemer G. Spermidine: a physiological autophagy inducer acting as an anti-aging vitamin in humans? Autophagy. 2019. doi:10.1080/15548627.2018.1530929

33. Niechcial A, Schwarzfischer M, Wawrzyniak M, Atrott K, Laimbacher A, Morsy Y, et al. Spermidine Ameliorates Colitis via Induction of Anti-Inflammatory Macrophages and Prevention of Intestinal Dysbiosis. J Crohns Colitis. 2023;17(9). doi:10.1093/ecco-jcc/jjad058

34. Jiang D, Guo Y, Niu C, Long S, Jiang Y, Wang Z, et al. Exploration of the Antioxidant Effect of Spermidine on the Ovary and Screening and Identification of Differentially Expressed Proteins. Int J Mol Sci. 2023;24(6). doi:10.3390/ijms24065793

35. Wang L, Jiang Y, Zhao C. The effects of advanced glycation end-products on skin and potential anti-glycation strategies. Experimental Dermatology. 2024. doi:10.1111/exd.15065

36. Li M, Sun J, Tan X. Evaluating the effectiveness of large language models in abstract screening: a comparative analysis. Syst Rev. 2024 Dec 1;13(1). doi:10.1186/s13643-024-02609-x PubMed PMID: 39169386.

